# Visualizing spatial population structure with estimated effective migration surfaces

**DOI:** 10.1101/011809

**Authors:** Desislava Petkova, John Novembre, Matthew Stephens

## Abstract

Genetic data often exhibit patterns that are broadly consistent with “isolation by distance” – a phenomenon where genetic similarity tends to decay with geographic distance. In a heterogeneous habitat, decay may occur more quickly in some regions than others: for example, barriers to gene flow can accelerate the genetic differentiation between groups located close in space. We use the concept of “effective migration” to model the relationship between genetics and geography: in this paradigm, effective migration is low in regions where genetic similarity decays quickly. We present a method to quantify and visualize variation in effective migration across the habitat, which can be used to identify potential barriers to gene flow, from geographically indexed large-scale genetic data. Our approach uses a population genetic model to relate underlying migration rates to expected pairwise genetic dissimilarities, and estimates migration rates by matching these expectations to the observed dissimilarities. We illustrate the potential and limitations of our method using simulations and geo-referenced genetic data from elephant, human and *Arabidopsis thaliana* populations. The resulting visualizations highlight important features of the spatial population structure that are difficult to discern using existing methods for summarizing genetic variation such as principal components analysis.

---

All natural populations exhibit some degree of “structure”: some individuals are more closely related than others. Population structure is shaped by many factors, but probably most influential are the barriers to gene flow that the population has experienced over its evolutionary history – barriers that may be due to extrinsic factors (such as topography or environment) or intrinsic factors (such as mate recognition, reproductive compatibility, or more complex interactions in social species such as humans). Studying the genetic structure of a population can therefore yield important scientific insights into the demographic and evolutionary processes that have shaped the population [1, 2], and help answer questions related to, for example, adaptation [3], speciation [4], hybridization [5], introgression [6] and recombination [7]. Understanding the genetic structure of a population may also be useful in contexts other than evolutionary genetics – for example, to identify subsets of genetically distinct groups that may require special conservation status [8], to detect the geographic origin of samples [9, 10, 11], or to help correct for confounding in genetic association studies [12, 13].

These important questions have motivated the development of many statistical methods for analyzing population structure. Among these, admixture-based clustering [14, 15, 16] and principal components analysis (PCA) [17, 18] are most widely used. A valuable feature of both approaches is that they summarize the main patterns of population structure in explicit and intuitive visual representations. Visual summaries are especially useful tools: not only can they help generate and refine hypotheses about the biological and evolutionary processes that have shaped the population, but they can also help identify sample outliers or other unexpected patterns, which are key steps in any analysis. Aside from this shared feature, admixture-based and PCA-based methods have their own distinct strengths and limitations. Clustering methods are particularly useful when the population under study is well represented by a small number of relatively distinct groups, possibly with recent admixture among groups. However, clustering methods are less well adapted to settings that exhibit more “continuous” patterns of genetic variation, such as “isolation by distance” where genetic similarity tends to decay with geographic distance [19]. In comparison, PCA is arguably better adapted to more continuous settings [17] and has proven helpful in diagnosing isolation by distance as a feature of the data [20]. However, certain properties of PCA complicate its interpretation and limit the insights it can provide. For example, PCA is heavily influenced by sampling biases: more data being collected preferentially from some regions than others [21, 22, 23]. And while PCA projections are often interpreted post hoc with geographic information in hand, PCA itself ignores the sampling locations even if they are known – information that can be particularly helpful if the data exhibit some degree of isolation by distance.

Motivated by this, we have developed a novel tool for visualizing population structure in an important setting that is not ideally served by existing methods: the setting where individuals are sampled from known locations across a spatial habitat (the samples are “geo-referenced”) and where the population structure is broadly, but perhaps not entirely, consistent with isolation by distance. We aim to produce visualizations which highlight regions that deviate from exact isolation by distance, and thus identify corridors or barriers to gene flow, if they exist. Our work shares goals with several previous methods [24, 25, 26], although the approach we take here is less algorithmic, as it explicitly represents genetic differentiation as a function of the migration rates in an underlying population genetic model. Our model-based approach is conceptually related to early work on inferring migration rates from genetic data [27], although the details – and particularly the use of “resistance distance”, a concept introduced in population genetics by [28] – are much closer to recent work on landscape connectivity [29] (see Discussion).

## Results

### Outline of the EEMS method

Figure 1 provides a schematic overview of our approach. In brief, the method is based on the “stepping stone” model [30], in which individuals migrate locally between subpopulations (“demes”), with symmetric migration rates that can vary by location. In order to capture “continuous” population structure, we use a dense regular grid of demes spread across the habitat, with each deme exchanging migrants only with its neighbors. Under the stepping stone model, expected genetic dissimilarities depend on the locations of samples and on the migration rates between connected demes. The expected genetic dissimilarity between a pair of individuals can be computed by integrating over all possible migration histories in their genetic ancestry, and we approximate it using a distance metric from circuit theory which integrates all possible migration routes between a pair of demes [28]. Our method effectively involves adjusting the migration rates so that the genetic differences *expected* under the model closely match the genetic differences *observed* in the data, while at the same time respecting the fact that nearby edges will often tend to have similar migration rates. The end result is an estimate of the migration rate on every edge in the graph, which we interpolate across the habitat to produce an “Estimated Effective Migration Surface” or EEMS. The EEMS provides a visual summary of the observed genetic dissimilarities among samples, and how they relate to geographic location. For example, if genetic similarities tend to decay faster with geographic distance in some parts of the space, this will be reflected by a lower value of the EEMS in those areas. If, on the other hand, genetic similarities tend to decay in the same way with distance throughout the habitat, the EEMS will be relatively constant. We use the term “effective” because the model makes assumptions – most importantly, equilibrium in time – that may preclude interpreting the EEMS as representing historical rates of gene flow. Nonetheless, as we illustrate on several examples, the method provides an intuitive and informative way to quantify and visualize patterns of population structure in geographically structured samples.

**Figure 1.**
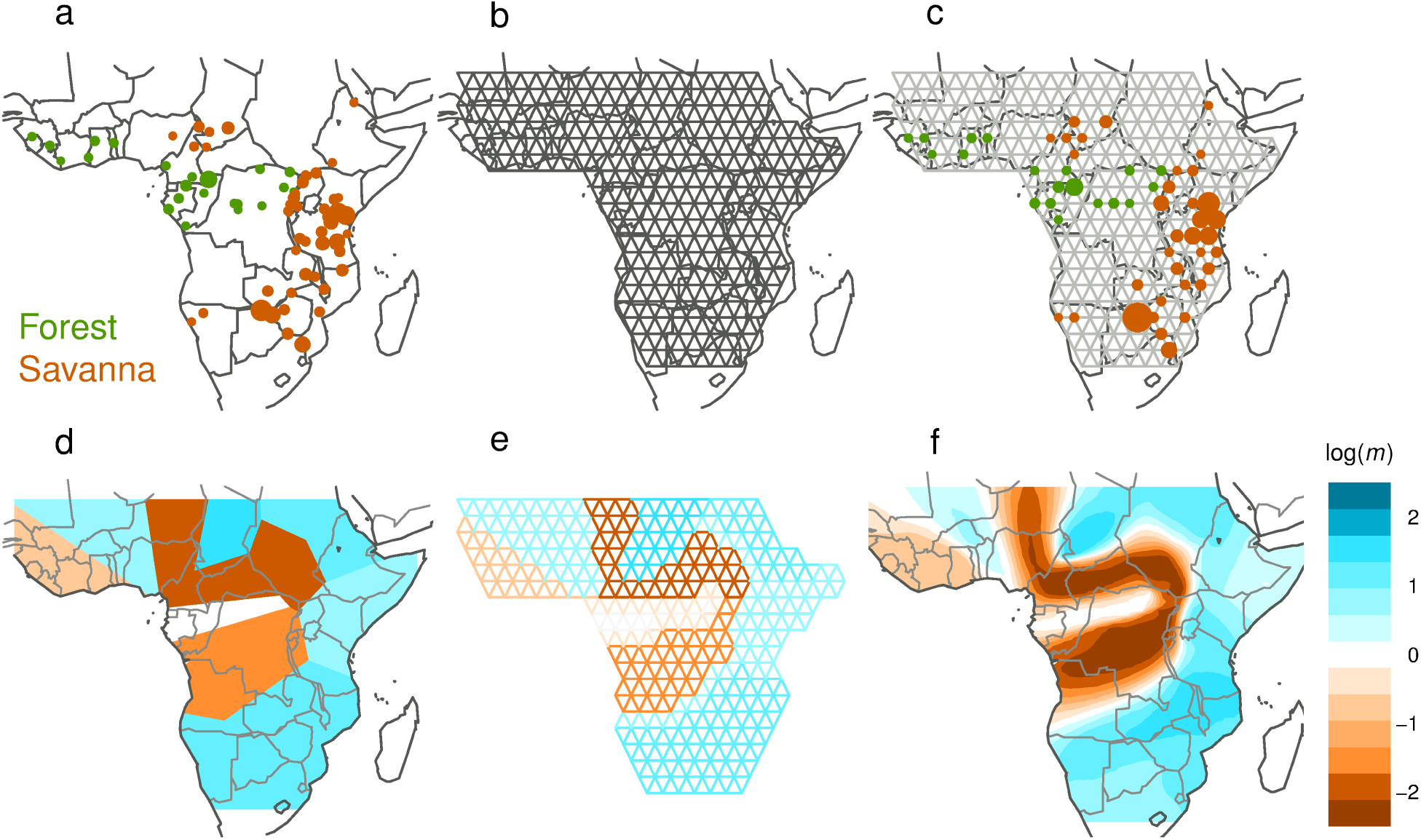
A schematic overview of the EEMS method, illustrated with geo-referenced African elephant data [32]. **(a-c)** Setting up the grid of demes: **(a)** Samples are collected at known locations across a two-dimensional habitat; green and orange colors represent two elephant subspecies – forest and savanna. **(b)** A dense triangular grid is chosen to span the habitat. **(c)** Each sample is matched to the closest deme on the grid. **(d-f)** Estimated Effective Migration Surface (EEMS) analysis: **(d)** Migration rates are allowed to vary according to a Voronoi tessellation which partitions the habitat into “cells” with constant migration rate; colors represent relative rates of migration in each cell, ranging from low (orange) to high (blue). **(e)** Each edge has the same migration rate as the cell it falls into. The cell locations and migration rates are then adjusted, using a Bayesian inference scheme, so that the expected genetic dissimilarities under the EEMS model closely match the observed genetic dissimilarities. **(f)** The EEMS is a colored contour plot, which is produced by averaging draws from the posterior distribution of the migration rates, interpolating between grid points. Here, and in all other figures, log(*m*) denotes the effective migration rate, on the log10 scale, relative to the overall migration rate across the habitat. (Thus log(*m*) = 1 corresponds to an effective migration rate that is 10-fold larger than the average.) The main feature of the elephant EEMS is a “barrier” of low effective migration that separates the habitats of the two subspecies: forest elephants to the west of the barrier, and savanna elephants to the north, south and east of the barrier.

### Simulations under the stepping stone model

We illustrate the benefits and limitations of the EEMS method with several simulations. Using the program ms [31], we simulated data from a stepping stone model under two different migration scenarios: a “uniform” scenario, in which migration rates do not vary throughout the habitat, intended to represent a pure isolation by distance situation (Fig. 2a); and a “barrier” scenario, in which a central region with lower migration rates separates the left and right sides of the habitat (Fig. 2b). We applied both EEMS, and – for comparison – PCA, to data generated under these scenarios and under three different sampling schemes (Fig. 2c). The results illustrate two key points. First, whatever the sampling scheme, the underlying simulation truth is much easier to discern from the EEMS contour plots (Fig. 2e) than from the PCA projections (Fig. 2d). For the pure isolation by distance setting, the EEMSs are approximately uniform under all three sampling schemes, and for the barrier scenario the EEMSs highlight the barrier as an area of lower effective migration. In contrast, the simple nature of the underlying structure is not obvious from the PCA projections for either scenario, and indeed, the PCA results for the different scenarios do not differ in an easily identifiable, systematic way. Second, EEMS is much less sensitive to the underlying sampling scheme than is PCA. Indeed, the inferred EEMSs are qualitatively unaffected by sampling scheme, except in the extreme case where there are no samples taken on one side of the migration barrier (which renders the migration rates on that side of the barrier inestimable from the data, so that estimates in that region are driven by the prior: no heterogeneity in migration rates). In contrast, PCA shows its known proclivity to be heavily influenced by irregular sampling [21, 22, 23]. For example, biased sampling and the presence of a barrier can both produce clusters in the PCA results (top row in Fig. 2d).

**Figure 2.**
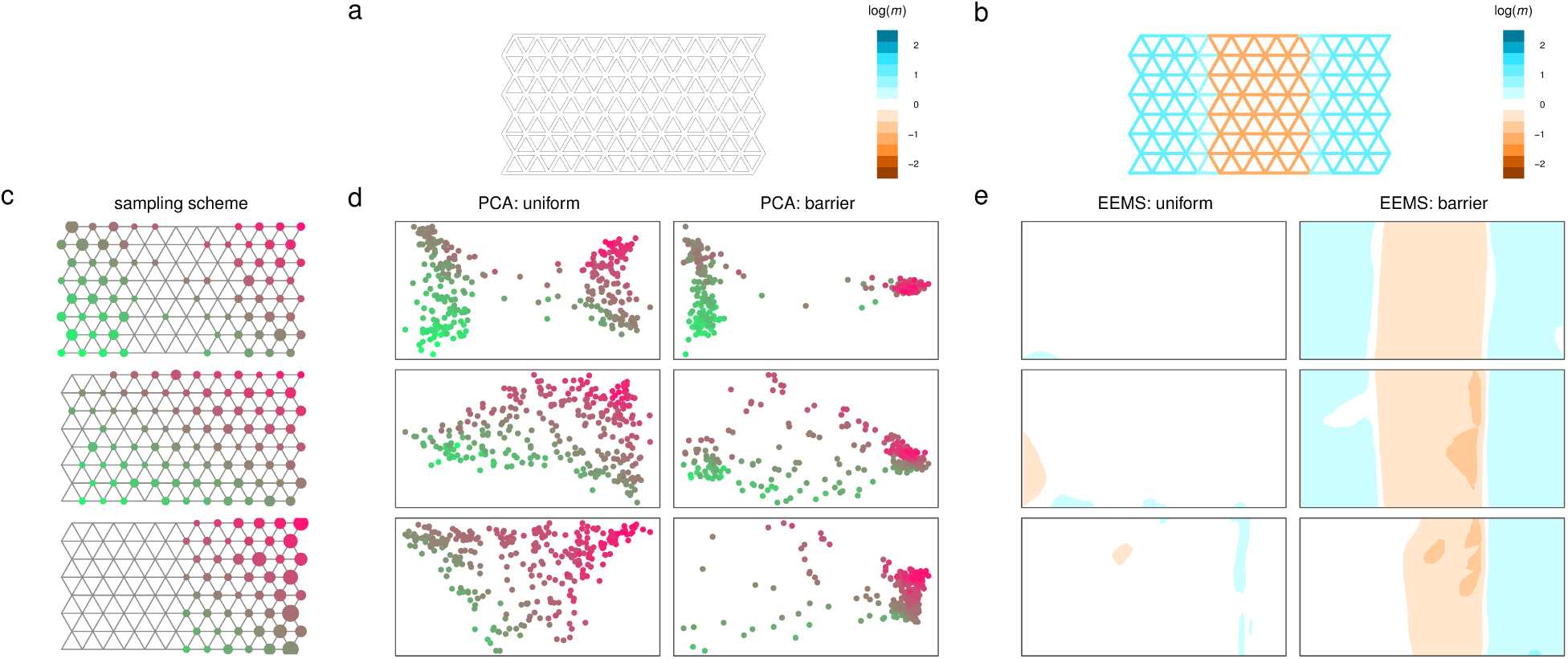
Simulations juxtapose EEMS and PCA analysis. For each method, we show results for two migration scenarios, representing “uniform” migration and a “barrier” to migration, and three different sampling schemes. **(a,b)** The true underlying migration rates under the two scenarios; colors represent relative migration rates. **(c)** The three sampling schemes used; the size of the circle at each node is proportional to the number of individuals sampled at that location, and locations are color-coded to facilitate cross-referencing the EEMS and PCA results. **(d)** PCA results. **(e)** EEMS results. In contrast to PCA, EEMS is robust to the sampling scheme and shows clear qualitative differences between the estimated effective migration rates under the two scenarios, which reflect the underlying simulation truth.

### Effective migration vs actual migration

Population genetics makes extensive use of the notion of “effective population size”, which can be informally defined as the size of an idealized (random-mating, constant-sized) population that would produce similar patterns of genetic variation as those observed in the population. The effective population size is typically quite different from the census size of the population. Similarly, the EEMS should be interpreted to represent a set of “effective migration rates” that, within an idealized stepping stone model evolving under equilibrium in time, would produce similar genetic dissimilarities as those observed in the data. Therefore, the effective migration rates will be different from actual migration rates of individuals in the population.

To illustrate this idea we present simulations under two different migration scenarios, each producing an EEMS with an “effective barrier” to migration, but for different reasons. In the first simulation the effective barrier results from a lower population density in the central region (Fig. 3a); in the second simulation the effective barrier results from the populations splitting some time in the past (Fig. 3b). In both cases the EEMS correctly reflects the structure in the observed genetic differentiation: individuals on either side of the central region are less genetically similar than expected based on distance alone (under pure isolation by distance). Indeed, in both cases the EEMS qualitatively reflects average rates of historical gene flow. However, it should be clear that care is warranted in linking the EEMS to inferences about the actual underlying migration processes.

**Figure 3.**
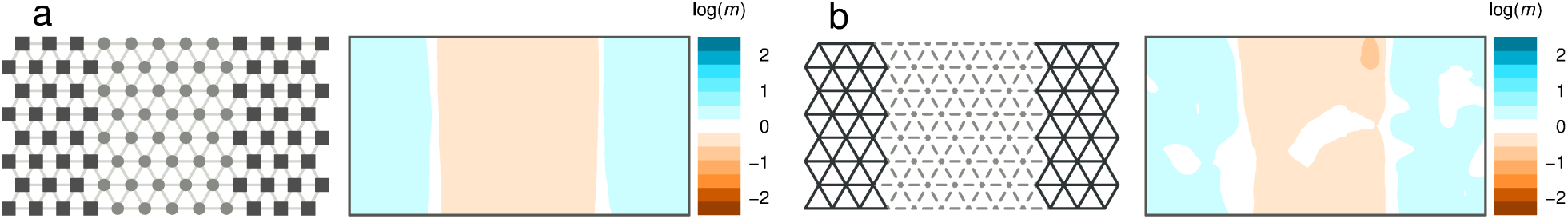
Simulations illustrate that EEMS infers effective migration rates, rather than actual steady-state migration rates. **(a)** Individuals have uniform migration rates but the central area has lower population density (those demes have fewer individuals, which is represented by smaller circles in gray). Thus fewer migrants are exchanged per generation. The difference in population density produces an effective barrier to gene flow, which is reflected in the EEMS. **(b)** A simple “population split” scenario: migration is initially uniform, but at some time in the past a complete barrier to migration arises in the central area (which is represented by dashed edges). Under this scenario the groups on either side of the central region diverge, which creates a barrier in the EEMS.

These results also illustrate the obvious but important fact that, because the EEMS method characterizes expected genetic differentiation under migration, it cannot distinguish between different scenarios that produce similar expectations for the pairwise genetic dissimilarities. This is also true of PCA [22]. In some cases it may be possible to distinguish among such scenarios using other aspects of the data, but we do not pursue this here.

### Effects of SNP ascertainment

Formally, the EEMS method, when applied to biallelic loci, assumes that these loci are “unascertained” – or, more precisely, that they are a random sample of biallelic loci polymorphic in the sample. This assumption would hold for complete sequence data, but not for SNPs on a genotyping chip, which are biased towards higher minor allele frequencies. Depending on the chip, SNPs could also be biased towards higher levels of polymorphism in certain geographic locations. To assess the robustness of EEMS to geographic bias in SNP ascertainment we performed simulations where SNPs were ascertained to be polymorphic in a small panel preferentially sampled from one geographic area. In these limited simulations, we found that the estimated effective migration surface is qualitatively robust to a biased ascertainment scheme (Supplementary Fig. 3a).

Although not a major focus of this paper, our method also estimates an effective diversity parameter within each deme, which reflects the expected genetic dissimilarity of two individuals sampled from that deme. In contrast to the effective migration rates, we found that the effective diversity rates are sensitive to geographically biased SNP ascertainment (Supplementary Figures 3b). This makes intuitive sense: the primary effect of ascertaining common SNPs is to increase the average number of differences among all individuals, increasing apparent diversity. Geographically biased ascertainment could therefore create apparent geographic differences in diversity where none exist. However, the effect of ascertainment on how, qualitatively, genetic dissimilarity *decays with distance* from any given location – and therefore its qualitative effect on our estimated migration surfaces – might be expected to be less pronounced, and our simulation results support this view.

### Anisotropic Migration

The model underlying EEMS assumes that migration rates are symmetric between pairs of adjacent demes, and so EEMS cannot infer a direction of migration on each edge. Nonetheless, EEMS is not entirely incapable of representing directional differences in migration (“anisotropic migration”). This is because, at any given deme, edges radiate in six directions, and each edge has its own migration rate. To illustrate, we simulated data where migration occurs at a much higher rate in the NS direction than in the EW direction: the resulting EEMS reflects this by interspersing vertical “corridors” that facilitate NS migration, with vertical “barriers” which inhibit EW migration (Supplementary Fig. 4).

### Diagnosing deviations from an EEMS fit

Although EEMS is formally based on a detailed demographic model – symmetric migration in a closed regular triangular grid, stationary in time – it would be a mistake to interpret the EEMS results as a validation of this specific model over other demographic models. Instead EEMS should be considered an exploratory tool for *representing and visualizing* patterns of genetic variation in geo-referenced data. In fact EEMS can provide useful representations of data that was not generated by the equilibrium stepping stone model (e.g., Figure 3b). However, not all patterns of genetic differentiation can be well represented by an EEMS. A general method to identify deviations from a model fit is to plot fitted values against observed values, and this is an approach we recommend here. Specifically, we suggest plotting the pairwise genetic differences predicted by the fitted EEMS against the pairwise genetic differences observed in the data. This plot can also be usefully compared with the analogous plot for a “pure isolation by distance” model (corresponding to constant effective migration) to identify aspects of the data that have been successfully captured by EEMS as well as aspects that have not. If a few individuals or sampling locations produce strong deviations from the EEMS fit, it may be prudent to remove those individuals before applying EEMS, or check that the results are robustness to their inclusion.

Supplementary Figure 5 illustrates these diagnostic plots for a situation where several individuals have recently migrated from one end of the habitat to the other (or perhaps had their sampling location wrongly labeled). The EEMS (Supplementary Fig. 5a) represents this with a barrier around the “migrants”, capturing the fact that they are genetically distinct from other near-by individuals. However, EEMS is unable to represent the fact that the “migrants” are genetically similar to some very distant individuals, as is evident in the diagnostic plot (Supplementary Fig. 5b). In principle this idea could be captured by a corridor of migration linking the “migrant” individuals to their original location, but EEMS does not do this, presumably because inserting such a corridor would make the overall model fit worse for other data points. We supply further examples of these diagnostic plots for all our empirical examples below (Supplementary Figures 11, 20 and 22).

## Empirical results

To illustrate EEMS in practice, we present results for four diverse empirical datasets: an African elephant dataset with strong differentiation between two geographically divided subspecies; two human datasets with individuals sampled from across Europe and across Sub-Saharan Africa where genetic differentiation has been shown to vary (somewhat) continuously with latitude and longitude; and an *Arabidopsis thaliana* dataset whose genetic variation is characterized by strong genetic similarity between Europe, where the plant is native, and North America, which it colonized in the last three hundred years [33].

### African elephants in Sub-Saharan Africa

The African elephant (*Loxodonta africana*) has two recognized subspecies – the forest elephant (*L. a. cyclotis*) and the savanna (or bush) elephant (*L. a. africana*). Both subspecies are under threat, partly from poaching, and a large sample was collected and genotyped at 16 microsatellite loci to help assign contraband tusks to their location of origin, and thus facilitate conservation efforts [9]. We analyze an augmented geo-referenced dataset from [32], which contains 211 forest and 913 savanna elephants.

The African elephant provides a helpful illustration of the EEMS method because the subspecies structure is clear and strongly correlated with geography, so we know the primary structure that we would like our method to highlight: the low effective gene flow between forest and savanna elephants despite their geographic proximity. And, indeed, the estimated effective migration surface is dominated by a strong barrier which seprates forest and savanna locations (Fig. 4b). Thus, the African elephant provides an empirical example of an effective barrier to migration due to a non-equilibrium history of drift after divergence, as in the simulation for Figure 3b. Notably, the EEMS successfully captures some of the winding shape of the geographic barrier between forest and savanna habitats, despite the fact that our method, based on Voronoi tessellations, seems better adapted to capture barriers with simpler structure.

**Figure 4.**
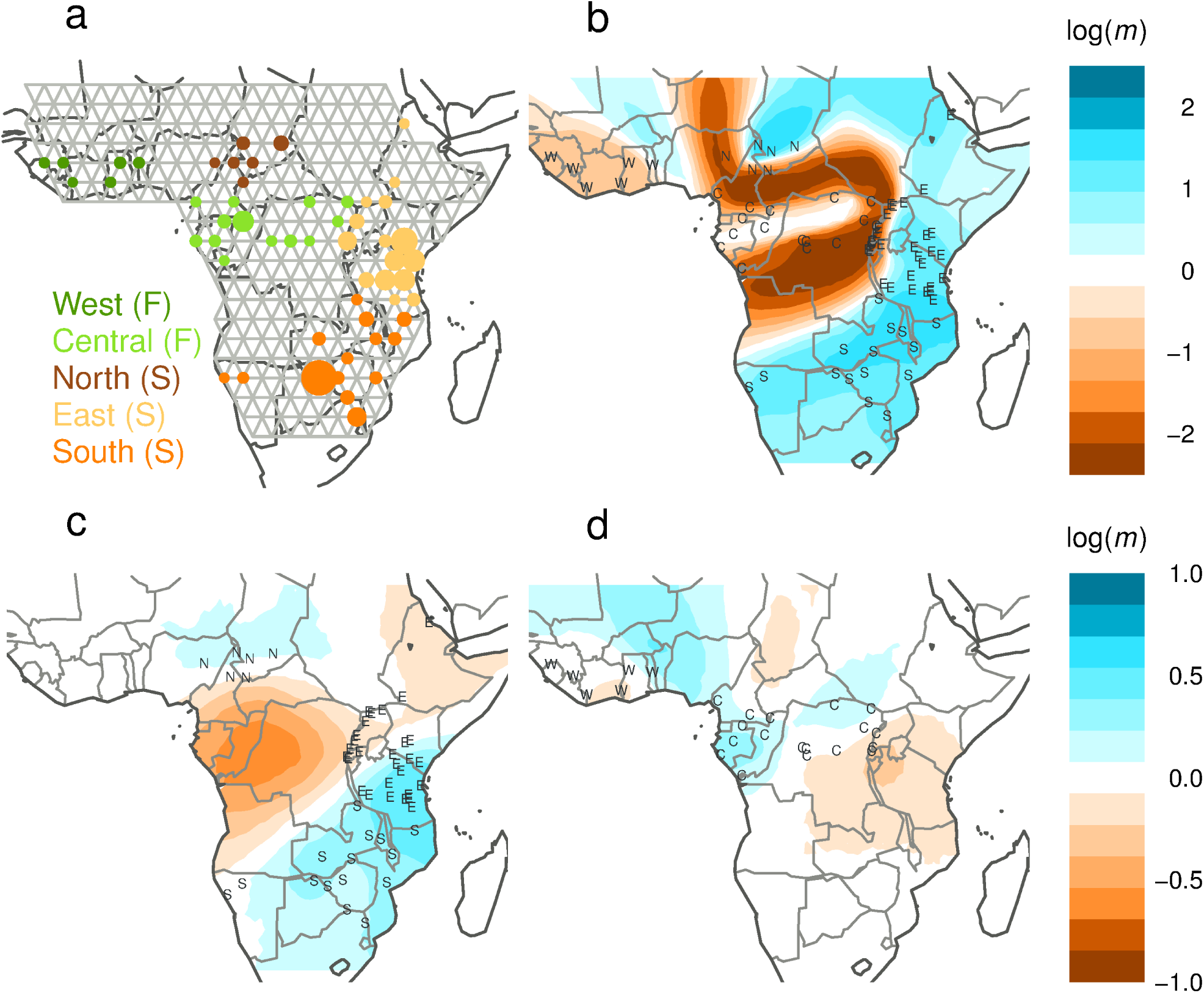
EEMS analysis of African elephant data from [32]. **(a)** African elephant samples are collected from two subspecies in five biogeographic regions: the forest elephant subspecies (in green) inhabits the west and central regions; the savanna elephant subspecies (in orange) inhabits the north, east and south regions. **(b)** Estimated effective migration rates for forest and savanna samples analyzed jointly. **(c,d)** Estimated effective migration rates for savanna and forest samples, respectively.

For the African elephant, one of the sixteen genotyped loci is extremely informative: the EEMS inferred from this locus alone is similar to that from all sixteen loci (Supplementary Fig. 9). However, the EEMS from the remaining fifteen loci is also similar to the EEMS for all sixteen loci (Supplementary Fig. 10a), demonstrating that this strongly differentiated locus is consistent with the others. (In principle, differences in inferred EEMSs among loci could provide a test for selection [34], but we do not pursue this here.)

Because the strong differentiation between forest and savanna elephants dominates the EEMS, we also analyzed forest and savanna samples separately to examine subtler structure within each group. The presence of substructure has been detected previously [9] as the African elephant habitat can be divided into five broad biogeographic regions: West and Central forest regions, and North, East and South savanna regions (Fig. 4a). The EEMS for savanna elephants (Fig. 4c) shows a barrier separating elephants in the North region from the rest, and a “corridor” of higher effective migration connecting the South and the East regions. The barrier coincides with forest habitat that forms a known barrier to migration for the savanna elephant; the corridor is consistent with previous observations, from mitochondrial data, that the South and East regions are genetically more similar than their geographic distance would suggest [35]. The EEMS for forest elephants (Fig. 4d) also shows a “corridor” of higher effective migration, which connects the West and the Central regions. Considered together, the two subspecies-specific EEMS plots suggest that there is more deviation from uniform migration (i.e., a stronger deviation from isolation by distance) in the savanna elephants. These patterns are harder to recognize in the corresponding PCA plots (Supplementary Fig. 6) or admixture and cluster-based analyses (Supplementary Figures 7 and 8).

In addition to the migration rates, our method also estimates an effective diversity parameter within each deme, which reflects the expected genetic dissimilarities of two individuals sampled from that location. For the African elephant, the inferred effective diversities are higher in the forest regions than in the savanna regions (Supplementary Fig. 10b). This represents in a direct, visual way the observation that forest elephants have higher heterozygosity than savanna elephants [36].

### Humans in Europe and Sub-Saharan Africa

We analyze two large-scale genome-wide datasets to visualize the genetic structure of human populations on two continents: a collection of 1201 individuals from 13 Western European countries genotyped at 197K SNPs [20, 37] and a collection of 314 individuals from 21 Sub-Saharan African ethnic groups genotyped at 28K SNPs [38].

Previous PCA-based analyses of both datasets [20, 38, 39] have found that the two leading PCs are correlated with geographic location. This suggests that genetic similarity tends to decay with geographic distance (Supplementary Fig. 13), and therefore, in broad terms, the data are consistent with isolation by distance. EEMS analysis on the other hand highlights patterns that deviate from *stationary* (exact) isolation by distance (Fig. 5). However, for both datasets, one must exercise caution with detailed interpretations of the EEMS results as the individual geographic locations are known only coarsely (see Discussion).

In Europe (Fig. 5a), the areas of highest effective migration span the North Sea and the Mediterranean, likely due to historic contacts between populations bordering these bodies of water; other areas of high migration span central France and Austria. Some regions of lower effective migration align with topographic barriers: the Alpes and the Atlantic. An area of low migration also spans Germany. Overall the results are consistent with the idea that population structure in Europe is characterized by a north/south cline [40], but the EEMS also elucidates more complex patterns of differentiation in the north/south direction. While the PCA plot may visually suggest a simple relationship between genetics and geography, the EEMS is far from a constant surface as one would expect under stationary isolation by distance. Thus the EEMS highlights patterns in the observed genetic dissimilarities that are difficult to discern in the corresponding PCA projections [20]. In POPRES, the geographic information is imprecise as sampling locations have been assigned based on nationality. However, the EEMS results are largely robust to the location uncertainty, which we assessed by adding random jitter to the assigned locations (Supplementary Fig. 16).

In Africa (Fig. 5b), the EEMS highlights a corridor of higher effective migration along the Atlantic coast, relative to lower effective migration inland. This indicates that – at a given distance apart – the coastal populations are more genetically similar than the inland populations. The U-shaped tail of the corridor moving inland suggests higher than expected (from the geographic distance alone) genetic similarity between some ethnic groups in the west and those in the east. Non-genetic information about the subpopulations can help clarify this pattern: in this case, the Fang (Fa) and the Kongo (Ko) in the west, and the Luhya (Lu) in the east speak Bantu languages, so we might hypothesize that the link is partly due to shared ancestry between Bantu speaking groups. In an EEMS analysis after excluding the Luhya, the definition of the corridor that connects the east with the west is greatly decreased (Supplementary Fig. 18), which supports the hypothesis that U-shaped signal is driven partly by the genetic similarity between Bantu speaking peoples (Supplementary Fig. 19).

**Figure 5.**
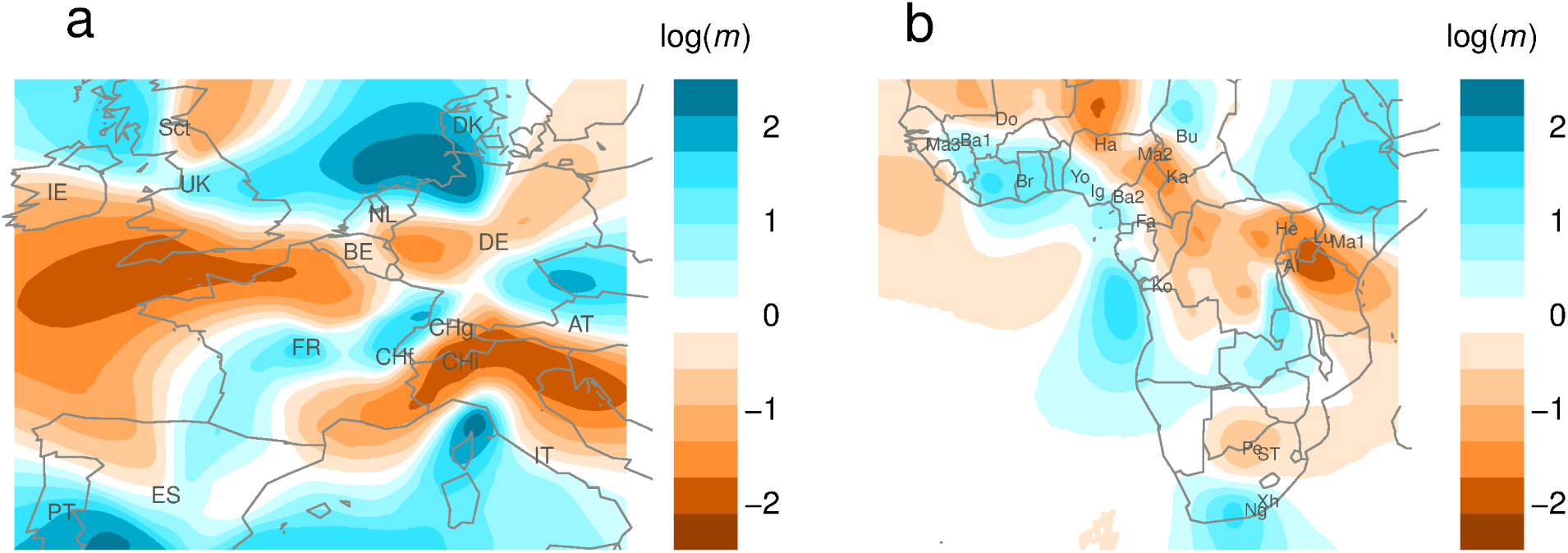
EEMS analysis of human population structure in Western Europe and in Sub-Saharan Africa. **(a)** Effective migration rates in Western Europe, estimated using geo-referenced data from the POPRES project [37]. **(b)** Effective migration rates in Sub-Saharan Africa, estimated using geo-referenced data from [41] and [42].

The EEMS method attempts to explain observed genetic dissimilarities using a stepping stone model with varying migration rates. Some datasets may contain features that are not captured by this model – for example, recent long-distance migrants. As a model-checking diagnostic we suggest comparing the fitted genetic dissimilarities and the observed genetic dissimilarities. For both human datasets, the fitted and observed dissimilarities generally agree well (Supplementary Figures 11 and 20). Furthermore, they agree much better than under a constant migration model: for example, the proportion of variance explained increases from 14.2% to 97.8% for the European data, and from 16.4% to 91.4% for the African data. This illustrates that the estimated non-stationary migration pattern is a better explanation for the observed patterns of spatial differentiation than exact isolation by distance.

### *Arabidopsis thaliana* in Europe and North America

*Arabidopsis thaliana* is a small flowering plant with natural range in Europe, Asia and North Africa, and which is now found in North America as well. Although *A. thaliana* is a selfing plant with low gene flow, its genetic variation has significant spatial structure [43, 44]. On the continental scale, in Europe the data exhibit patterns consistent with isolation by distance, with an east/west gradient that has been interpreted as evidence for post-glaciation colonization [43]. In North America there is less spatial structure, genome-wide linkage disequilibrium and haplotype sharing, likely due to recent human introduction from Europe [43].

We analyze a large geo-referenced dataset from the Regional Mapping (RegMap) project [45]. The data include 980 accessions from Europe and 180 accessions from North America, genotyped at 220K SNPs.

In a combined analysis of the North American and European data (Fig. 6a), the EEMS shows a corridor of high effective migration across the Atlantic Ocean, relative to lower effective migration within each continental group. This highlights the fact that the European and North American samples are more genetically similar than their distance would suggest under a simple isolation by distance scenario. Although the EEMS model assumes that migration is symmetric, and so it cannot infer a direction for gene flow, the EEMS is consistent with the hypothesis that recent directed migration introduced *A. thaliana* from Europe to North America.

Analyzing the North American samples alone, the EEMS has an area of high migration connecting the two sampled regions, Lake Michigan and the Atlantic coast (Fig. 6b). This indicates that samples from these two regions are similar genetically even though they are distant geographically – probably again due to human-assisted long-range “migration” rather than natural dispersal, and consistent with the observation that there is extensive haplotype sharing not only within but also between sampling locations [43].

**Figure 6.**
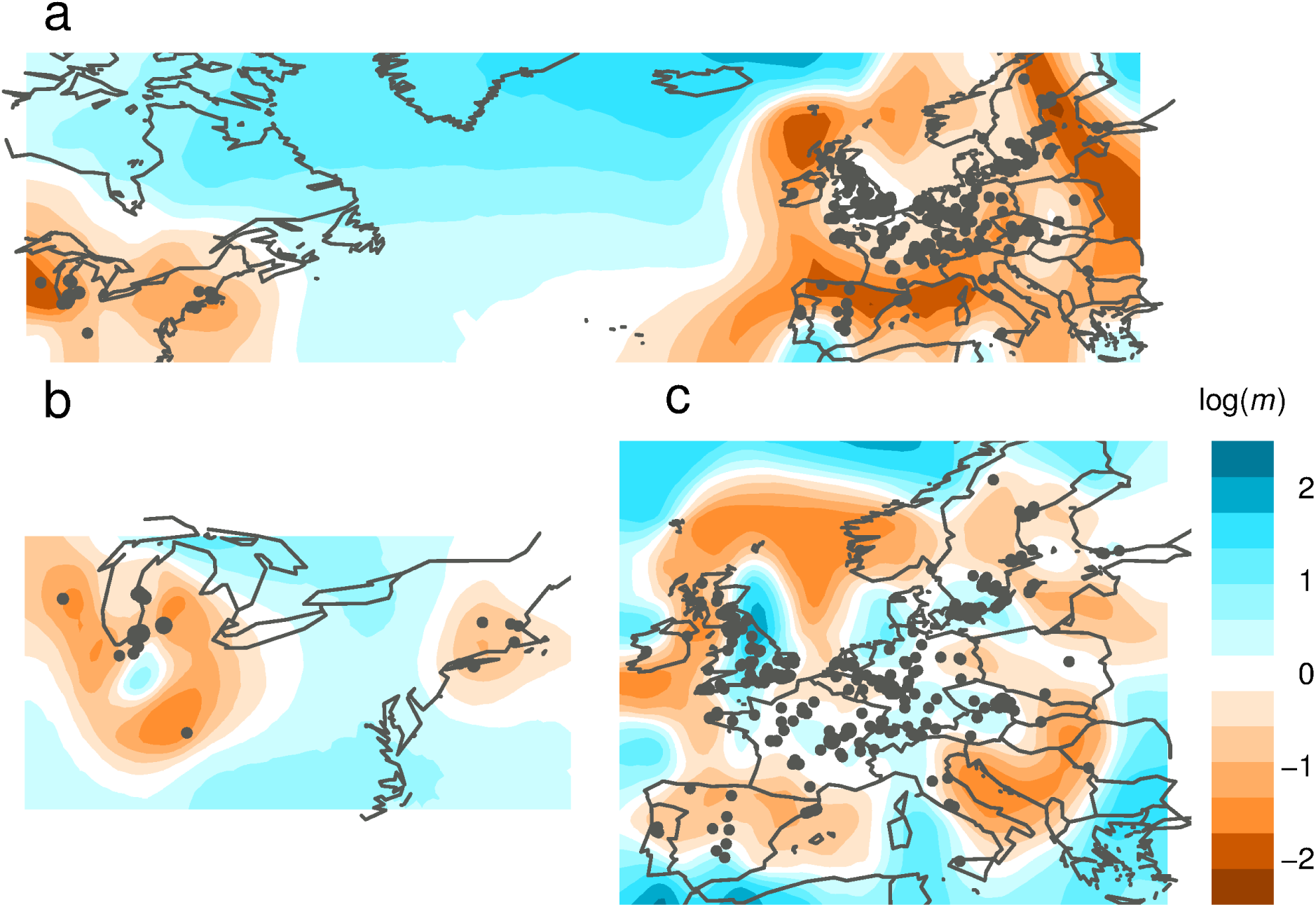
EEMS analysis of *Arabidopsis thaliana* data from the RegMap project [45]: **(a)** Estimated effective migration rates in North America and Europe combined; **(b)** Estimated effective migration rates in North America; **(c)** Estimated effective migration rates in Europe.

For the European samples the EEMS highlights a number of regions of higher and lower effective migration (Fig. 6c). For example, in the British Isles, a region of lower migration separates the northern British Isles from the rest of Britain, which in turn is connected to NW France by an area of higher migration. (In PCA analysis in [45] most accessions from the British Isles are projected closest to France.) The lower effective migration area in Central and Northern France highlights the fact that the NW France samples are more similar to British samples than they are to other French samples a similar distance away. An area of higher effective migration covers much of Germany, and links it with eastern France in the south, and to parts of Norway and Sweden in the north. In contrast, Spain and Italy show substantially lower effective migration rates, suggesting that genetic similarity decays more quickly with distance in these areas than in other parts of Europe.

## Discussion

We have developed EEMS (Estimated effective Migration Surfaces), a new method for analyzing population structure from geo-referenced genetic samples. Our method produces an intuitive visual representation of the underlying spatial structure in genetic variation, which highlights potential regions of higher-than-average and lower-than-average historic gene flow. EEMS is specifically applicable to data that conform roughly to “isolation by distance” (IBD), i.e., in settings where genetic similarity tends to decay with geographic distance, but where this decay with distance may occur more quickly in some regions than in others.

EEMS uses the concept of “isolation by resistance” (IBR), which aims to model how genetic differentiation accumulates in non-homogeneous landscapes [28]. IBR integrates over all possible migration paths between two points, providing a computationally convenient approximation to the coalescent process in structured populations, and better prediction of genetic differentiation than alternatives such as the least-cost path distance [46].

As originally introduced, IBR is used to build up a resistance map from known landscape/habitat features and thus determine whether resistance distance can improve prediction of observed genetic differentiation over Euclidean or least-cost path distances [46]. More recently, IBR has been incorporated into a formal inference procedure designed to test whether resistance distances are correlated with specific, known landscape features such as altitude or river barriers [29]. In contrast, EEMS estimates an effective migration surface from genetic data without the need to observe underlying environmental variables, and thus provides an exploratory tool for spatial population structure. We view the hypothesis-driven and exploratory approaches as complementary, and we anticipate that each could be useful in many applications.

EEMS also complements existing methods for detecting barriers to gene flow in continuous spaces, such as Monmonier’s maximum difference algorithm [24], Wombling methods [25] and LocalDiff [26]. Monmonier’s algorithm analyzes observed genetic differences, but it requires that genetically distinct groups are specified *a priori* before attempting to find strongly differentiated pairs. Wombling methods can detect strong barriers without pre-specified clustering, by estimating spatial gradients in allele frequencies and identifying localized sharp discontinuities. LocalDiff [26] uses a spatial Gaussian process to interpolate allele frequencies before assessing local patterns of differentiation among neighboring populations. The algorithmic flavor of these methods contrasts with EEMS, which directly models landscape inhomogeneity and uses isolation by resistance to do so.

Although EEMS is aimed at settings where the underlying population structure is somewhat *continuous*, our method is built on a dense regular grid of *discrete* demes, with migration between neighboring demes. Since the demes do not correspond to *a priori* defined subpopulations, the size and registration of the grid are somewhat arbitrary. The choice of grid may be influenced by factors such as the density of sampling locations (one might want a grid sufficiently dense so that different sampling locations typically correspond to different demes), and computational tractability (computation scales cubically with number of demes in the grid). Although we have found that in practice results are qualitatively robust across a range of grids (Supplementary Fig. 1), some details can change with the specific grid, and so we suggest reducing the potential influence of the grid choice by averaging results over several different grids. In principle, it would seem attractive to dispense with the grid altogether, and use models of continuous migration. However, such models present theoretical and mathematical challenges [47], and at present we do not know how to achieve this.

In addition to estimating effective migration rates, our model also fits a deme-specific parameter, *q*, which describes the within-deme genetic diversity. For the purpose of characterizing dissimilarities between demes, *q* is a nuisance parameter, but in some applications spatial variation in diversity may be of interest in itself, and so visualizing the effective diversity rates may be useful for some analyses. In our examples, visualizing *q* highlights the previously noted north-south gradient in diversity in human Europeans (Supplementary Fig. 14b) higher diversity in ethnic groups from East Africa compared to groups from South or West Africa (Supplementary Fig. 17b), and higher diversity in forest versus savanna elephants (Supplementary Fig. 10b). EEMS requires that each sample has a specified geographic origin, but in some cases this information may be known only imprecisely. For example, in our analysis of human genetic variation in Europe, we use the coordinates previously described in [20], where samples are assigned to the geographic center of their ancestral country of origin, with the exception of samples from the Russian Federation, Sweden and Norway, which are assigned to the capital. To better deal with imprecision in geographic origin, the uncertainty could be incorporated into the model, as in [15]: the actual location of each individual will be treated as an unobserved latent variable, given a prior distribution, and integrated out in the MCMC estimation scheme. This approach might also improve robustness to data errors such as sample switches, and identify individuals whose genetic origins differ appreciably from their physical sampling locations. A similar extension could also help address the “spatial assignment problem” in non-stationary isolation by distance settings, i.e., attempt to infer the origin of individuals with unknown location, given reference samples with known location [9, 10].

Since the primary output of EEMS is a visual display of spatial patterns, we have paid attention to the details of this display. For example, we have selected a color scheme that is colorblind friendly [48] and that is “balanced” with respect to high versus low migration. (For example, we have attempted to give similar visual prominence to regions that are 10 times higher than average in their effective migration as to regions that are 10 times lower than average.) We have chosen the scale so that small differences in effective migration rates say, less than a factor of two – tend not to be emphasized. These choices could undoubtedly be improved upon with further experimentation, and indeed any given scale or color scheme may work better for some datasets than for others. Users of the EEMS method may therefore wish to experiment with display settings, and should be aware of the impact that display parameters can have on the message conveyed by an EEMS.

Although we have not emphasized it here, in addition to a point estimate of the effective migration surface, EEMS fits a posterior distribution for the migration parameters at each location. Given that our modeling assumptions are simplistic we would not advocate interpreting these posterior distributions too literally; nonetheless, they may provide a useful assessment of the uncertainty in the estimated surface. It could be helpful to incorporate this information into the visual display: for example, shading the EEMS only in regions where the posterior 95% credible interval excludes 0 could reduce the danger of over-interpreting patterns that could have easily arisen by chance.

Regarding computation time, for the analyses presented here we have used population grids that range from 120 to 520 vertices. The current implementation does not scale well beyond 1000 vertices because the computational cost is cubic in the size of the grid. We typically run the MCMC sampler for at least 8 million iterations, which takes about 15 hours of actual CPU time for a grid with 500 demes. However, for assessing convergence, it is important to simulate several realizations of the Markov chain, i.e., starting the EEMS program several times with a different random seed. For the analyses presented here we have averaged results across at least 8 independent realizations. The software also allows restarting the MCMC sampler if the diagnostic posterior trace plot indicates the Markov chain has not converged in the specified number of iterations.

Like PCA, EEMS works directly with a matrix that summarizes the (dis)similarities between all pairs of samples, by averaging across genotyped markers. This makes it computationally tractable for large numbers of SNPs: once this matrix is computed, the complexity per MCMC iteration does not depend on the number of markers. Moreover, like PCA, EEMS is widely applicable: it can be applied to visualize dissimilarities between geo-referenced samples that have been computed from non-genetic features, e.g., language. Summarizing the data by a pairwise distance matrix does, however, result in some loss of information. In particular, as highlighted in Figure 3, it limits the demographic scenarios that can be distinguished from the data; see [22] for detailed discussion. It may be fruitful to explore the use of other dissimilarity matrices to emphasize different aspects of the data, perhaps different historical timescales. For example, ChromoPainter [49] produces a measure of genetic similarity (“number of chunks copied”), which will tend to emphasize the most recent coalescent events between samples (rather than the average coalescence times). Similarly, distance matrices based on rarer SNPs should tend to emphasize more recent dispersal history [50]. It is possible that the pairwise sequentially Markov coalescent [51], extended to deal with pairs of individuals [52], could also provide useful alternative distance matrices that could be visualized as an EEMS.

Software implementing the EEMS method is available at http://www.github.com/dipetkov/eems.

## Acknowledgements

This work was supported in part by US National Institutes of Health (NIH) grants CA198933 (J.N.,M.S.) and grant HG02585 (M.S). We thank Samuel Wasser for access to the African elephant data from [32] and Ida Moltke for compiling the human dataset from Sub-Saharan Africa as described in [38]. We also acknowledge Brad McRae for helpful early discussions on resistance distances.

## Online methods

EEMS uses a population genetic model that involves migration on an undirected graph *G* = (*V*, *E*) with vertices (demes) *V* connected by edges *E*. The graph *G* is a regular triangular grid, which is fixed and embedded in a two-dimensional plane, so that each deme has a known location and only neighboring demes are directly connected (Figure 1b). The density of the grid is pre-specified by the user, and depends on both computational considerations – computational complexity scales cubically with the number of vertices – and the resolution of the available spatial data.

The EEMS model has migration parameters *m* and diversity parameters *q* such that *m* = {*m*_*e*_ : *e* ∈ *E*} specifies an effective migration rate on every edge in *E* and *q* = {*q*_*v*_ : *v* ∈ *V*} specifies an effective diversity rate for every deme in *V*. Intuitively, the migration rates *m* characterize the genetic dissimilarities between distinct demes, while the diversity rates *q* characterize the genetic dissimilarities between distinct individuals from the same location. The EEMS model is a particular application of the more general stepping stone model [30], which allows directed migration as well as migration between demes that are not located close in space.

We use Bayesian inference to estimate the EEMS parameters *m* and *q*. The key components to this inference are the *likelihood* 𝓁(*m*, *q*), which measures how well the parameters explain the observed data, and the *prior distribution p*(*m*, *q*), which captures the expectation of a spatial structure in the parameters *m* and *q*. In particular, the prior captures the idea that the migration rates on adjacent edges of the graph are often similar. The following sections describe the likelihood and the prior in detail.

### The likelihood

We first specify the likelihood for SNP data (on *n* individuals at *p* SNPs), and then extend it to microsatellites. The key initial step is to summarize the observed genetic data by the matrix of average genetic differences, *D*, between every pair of sampled individuals. (The matrix *D* defined precisely below.) This approach – using the matrix of pairwise dissimilarities as a sufficient statistic for the population parameters – is motivated by the assumption that *D* contains most of the information about *m* and *q*. This may not be completely true, but the idea of performing inference using pairwise genetic similarities or dissimilarities has a long history in both population genetics and phylogenetics [53, 54, 55], and many existing methods are based on a similar assumption. For example, PCA [17] and TreeMix [56] both work with the genetic covariance matrix.

Therefore, let *D*_*ij*_ denote the observed genetic dissimilarity between individuals *i* and *j*. The expected value of *D*_*ij*_ is determined, up to a constant of proportionality that reflects the mutation rate, by how closely related *i* and *j* are, or more precisely, by the expected coalescence time between their gametes. The pairwise expected coalescence time in turn depends on the sampling locations δ(*i*), δ(*j*) and on the population parameters (*m*, *q*): individuals sampled from demes that are connected by many short paths containing edges with high migration rates will tend to be more closely related, and hence more similar genetically, than individuals sampled from demes connected only by paths that are long and/or contain edges with low migration rates. The expected coalescence times can be computed, at some computational expense, by solving a large set of simultaneous equations; alternatively, they can be approximated – at less, but still nontrivial computational cost – using the concept of “resistance distance” [28]. We implemented both approaches, and found them to produce qualitatively similar effective migration surfaces, and so all results presented here were obtained using resistance distances.

Letting *σ*^2^ denote the constant of proportionality mentioned above, we can write:

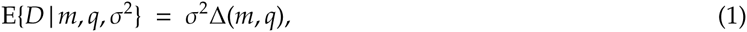

where Δ(*m*, *q*) is a matrix of expected dissimilarities that can be computed for any *m* and *q*. Our modeling approach, detailed below, assigns high likelihood to values for (*m*, *q*, *σ*^2^) such that *σ*^2^Δ(*m*, *q*) ≈ *D*, while taking some account of dependencies among elements of *D* and of linkage disequilibrium among markers.

To make our specification precise, we introduce the following notation:

- *Z* is the *n* × *p* matrix of genotypes: *Z*_*il*_ is the genotype of individual *i* at locus *l*, which is coded as 0, 1 or 2 copies of the minor allele.
- *D* is the *n* × *n* matrix of average genetic differences between individuals: 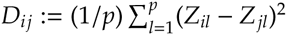.
- *L* is the (*n* - 1) × *n* matrix such that *L*_*i*_ = *e*_*i*_ - *e*_*i*+1_ where *L*_*i*_ is the *i*th row of *L* and *e*_*i*_ is a row vector with 1 in the *i*th component and 0s elsewhere.
- *W* is the (*n* - 1) × (*n* - 1) matrix -*LDL*′ = 2(*LZ*)(*LZ*)′, which is positive definite as a quadratic form [29].

The matrix *L* is chosen because it forms a basis for the space of contrasts on *n* elements: for example, *e*_*i*_ - *e*_*i*+1_ is a contrast between the *i*th and (*i* + 1)st elements. Since *L* is a basis, *W* is a one-to-one mapping of *D* and we can specify a model for *D* by specifying a model for *W* [29]. It turns out that, for technical reasons, it is easier to work with *W* than with *D* directly [57]. Furthermore, since the statistic *W* is positive definite, a natural model for *W* is the Wishart distribution, which can be parametrized by the expectation, E{*W*}, and a scalar degrees of freedom parameter, *k*. Using equation (1), the expectation is given by:

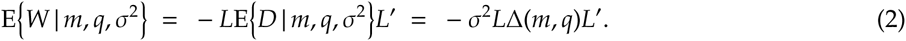

We treat the degrees of freedom *k* as an additional free parameter to be estimated.

Putting this together yields a closed form for the density of *W* and thus – for the likelihood of the parameters *k*, *m*, *q*, *σ*^2^ since 𝓁(*k*, *m*, *q*, *σ*^2^) := *p*(*W* | *k*, *m*, *q*, *σ*^2^). Specifically:

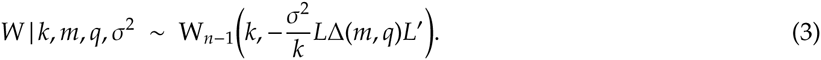

We make the following observations:

1. Although we defined *Z*_*il*_ as the number of copies of the minor allele, the likelihood does not depend on the labeling of the alleles because the differences (*Z*_*il*_ - *Z*_*jl*_)^2^ do not depend on the labeling.
2. If the genotypes *Z* were independent across loci and normally distributed, then standard Gaussian theory would imply that *W* has a Wishart distribution, *with the degrees of freedom k equal to the number of SNPs p*. However, genotypes are neither normal nor independent, and rather than fix *k* = *p* (as in [29]), we estimate the degrees of freedom *k*, under the assumption *n* ≤ *k* ≤ *p*. The smaller *k* is, the higher the variance of *W* about its expectation. (E{*W*} does not depend on *k* in our parametrization.) By allowing *k* < *p* we can, to some extent, account for sources of model mis-specification such as linkage disequilibrium between loci.
3. In defining *W* := -*LDL*′ we took *L* to be a specific matrix. However, other choices for *L* would yield equivalent likelihoods as long as the (*n* - 1) rows of *L* form a basis for the contrasts of *n* elements. (A contrast is a linear combination whose coefficients add to 0.) This key property ensures that *W* is a one-to-one mapping of *D*, and therefore we would get exactly the same likelihood with any basis *L* (up to a constant of proportionality that does not depend on the parameters). In other words, the introduction of a specific transformation *W* should be regarded as a technical trick to facilitate the development of a model for the genetic differences *D*; see [57] for a more mathematical discussion.

### Application to microsatellites

At a microsatellite locus, an allele is typically coded as the number of repeats of a specific motif. To apply our method to microsatellites we define the genotype *Z*_*il*_ to be the *average* of the two alleles individual *i* carries at locus *l*. (This approach could likely be improved upon, but it suffices for our analysis of the African elephant data.)

We then define *D*^(*l*)^ as the matrix of pairwise differences at locus *l*, and *W*^(*l*)^ as the corresponding positive definite transformation:

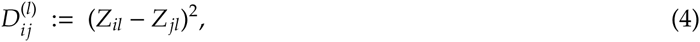

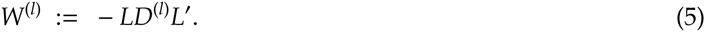

Since different microsatellite loci have different mutation rates, we introduce locus-specific scale parameters 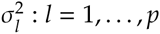. For locus *l* equation (1) becomes:

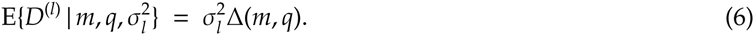

And we define the likelihood by assuming that each *W*^(*l*)^ has an independent Wishart distribution, with one degree of freedom:

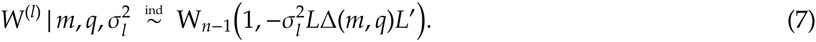

Each matrix *W*^(*l*)^ has rank one and hence is not positive definite but we can nevertheless compute the likelihood for the parameters *m*, *q* and 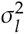. (Since the *p* microsatellite loci are assumed to be independent, the degrees of freedom are effectively fixed to *k* = *p*.)

### The dissimilarity matrix Δ(*m*, *q*)

In population genetics, the expected genetic dissimilarity between two samples is a function of their expected coalescence time. Indeed, for haploid samples at biallelic loci, as the mutation rate tends to 0, it can be shown that (see Supplementary Methods and [22]):

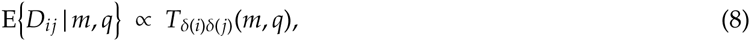

where δ(*i*) denotes the deme from which sample *i* is drawn, and *T*_αβ_(*m*, *q*) is the expected coalescence time of two independent haploid samples taken from demes *α* and *β*, respectively. Thus in equation (1) we have:

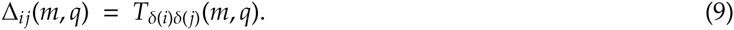

Similarly, for diploid samples, we have (see Supplementary Methods):

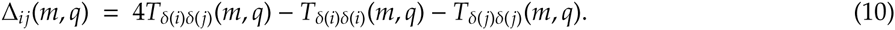

For any particular value of the parameters *m* and *q*, the matrix *T*(*m*, *q*) – and hence Δ(*m*, *q*) – can be computed by solving a system of linear equations [58, 59]. (In the stepping stone model *T*(*m*\*, *q*\*) is a function of he between-deme migration rates *m*\* and the within-deme coalescence rates *q*\*, where we use the superscript * to indicate that the EEMS parameters have slightly different interpretation from those in [58, 59].)

However, computing the matrix of expected coalescence times *T* is expensive because it requires solving a linear system with *d*(*d* +1)/2 unknowns to find all pairwise expected coalescence times in a graph with *d* demes – this has complexity O((*d*^2^)^3^). To reduce the computational cost, for all results presented here, we approximate coalescence times using the idea of “effective resistances” – a distance metric for weighted undirected graphs [60]. Computing the matrix of effective resistances *R* is less intensive because we can obtain all pairwise resistance distances by inverting a *d* × *d* matrix [61] – this has complexity O(*d*^3^). (Efficiency can be improved further by computing the subset of resistance distances between sampled demes only; see Supplementary Methods.)

To approximate coalescence times in terms of effective resistances, let *R*_αβ_(*m*) denote the resistance distance between demes *α* and *β* in the graph *G*. (Note that *R*_αβ_ is not a function of only the *local* migration rate *m*_αβ_, but is determined by the *global* migration pattern *m*.) The effective resistances *R* are approximately related to the expected coalescence times *T* through [28]:

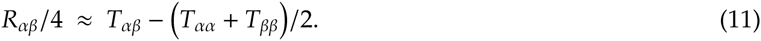

This approximation is exact for isotropic migration (i.e., if the demes are equivalent with respect to the rate and pattern of migration), and for more general migration models the approximation gets better as the migration rates increase [28]. Using equation (11) we approximate the expected coalescence time between two haploid samples from demes α and β as:

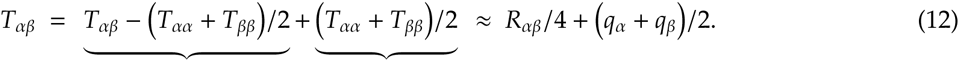

That is, for each pair of demes (α, β) w e split the ex pecte d coale scence time *T*_αβ_ into a between-demes component, which is approximated by the (scaled) effective resistance *R*_αβ_, and a within-deme component (*q*_α_ + *q*_β_)/2, which is determined by the diversity rates *q*. The effective resistances *R* = (*R*_αβ_) depend on *m*; the vector *q* is treated as a free parameter to be estimated. We then obtain Δ(*m*, *q*) by substituting *T*(*m*, *q*) with its approximation according to equation (12).

Generally, we have found that the approximation based on the effective resistances *R* produces visualizations comparable with those obtained using the more computationally expensive coalescence times *T*.

## Prior Distributions

### The Voronoi prior on migration rates

Our prior for the migration rates *m* captures the idea that nearby edges will tend to have similar rates, while it also allows the rates to vary among edges. We parametrize the prior *p*(*m*) using a Voronoi tessellation of the two-dimensional habitat *H*, which partitions *H* into *C* convex polygons (cells) as follows. First select *C* distinct points (seeds) *s*_1_, …, *s*_*C*_ within *H*. Then define cell *c* to be the set of points in *H* that are closer to seed *s*_*c*_ than to any other seed. Given a Voronoi tessellation of *H*, we associate with each cell a migration rate *m*_*c*_. We use these to induce a migration rate on each edge in the graph *G*, with the migration rate of the edge joining demes α and β given by:

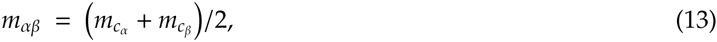

where *c*_α_ denotes the cell that contains deme α. Migration rates are naturally positive and therefore we parametrize the *m*_*c*_ values on the log10 scale. Further, to capture the idea that the migration rates of different cells may be similar to one another we parametrize them as deviations from an overall mean rate *μ*:

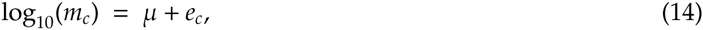

where the “effect” of cell *c*, denoted by *e*_*c*_, determines whether the local dispersal in cell *c* is faster or slower than the average.

In this formulation, migration rates on every edge in the graph are determined by the parameters (*C*, *s*_1_, …, *s*_*C*_, *e*_1_, …, *e*_*C*_, *μ*). To complete the Bayesian specification we place priors on the model parameters:

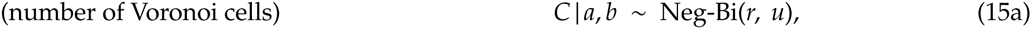

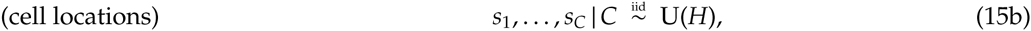

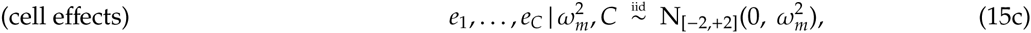

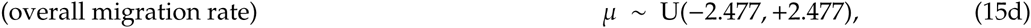

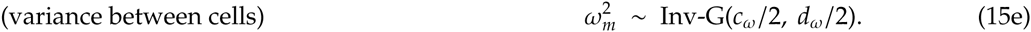

Here U(*H*) denotes the uniform distribution with support the habitat *H*; N_[*a*,*b*]_(*μ*, *σ*^2^) denotes the truncated normal distribution with mean *μ*, variance *σ*^2^ and support [*a*, *b*]; Inv-G(*c*, *d*) denotes the inverse gamma distribution with shape *c* and scale *d*; and Neg-Bi(*r*, *u*) denotes a zero-truncated negative binomial distribution with shape (number of failures) *r* and probability of success *u*. [The zero-truncated negative binomial has support {1, 2, 3, …}; we truncate the support at zero because the Voronoi tessellation should have at least one cell.] For all results reported here we have used *r* = 10, *u* = 2/3, *c*_*ω*_ = 0.001, *d*_*ω*_ = 1.

The prior on the number of Voronoi cells *C* was chosen because the negative binomial is an overdispersed Poisson (a continuous mixture of Poisson distributions): Suppose that *C* is a Poisson random variable with mean *λ*, and *λ* is a Gamma random variable with shape *r* and rate *u*/(1 - *u*). If we integrate with respect to ?, we obtain *C* ∼ Neg-Bi(*r*, *u*). For all analyses described here, we have used *r* = 10 and *u* = 2/3, which results in a diffuse prior on *C*, with prior mean 20 and prior variance 60.

The lower and upper bounds on the mean log migration rate *μ* are chosen so that on the original scale the mean migration rate varies in the range [1/300, 300]. The bounds are somewhat arbitrary, and chosen to reflect values that might be considered “very small” (approaching the limit of discrete demes evolving independently) and “very large” (approaching a panmictic population). The cell effects *e*_1_, …, *e*_*C*_ are constrained to lie in the range [-2, +2], so that the migration rate of a cell can vary within a factor of 100 from the mean migration rate.

### Other priors

If there are more genotyped markers than sampled individuals, we can estimate the degrees of freedom *k*. The prior on *k* is uniform on the log10 scale, to reflect our uncertainty about the order of magnitude of this parameter:

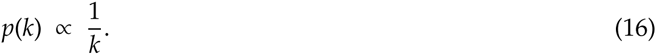

The prior is proper because *k* is bounded: *n* ≤ *k* ≤ *p* where *n* is the number of samples and *p* is the number of SNPs.

The prior on the Wishart scale parameter *σ*^2^ is:

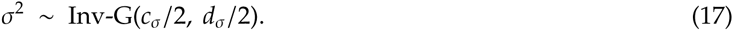

In all results presented here we have used *c*_*σ*_ = 0.001, *d*_*σ*_ = 1.

### General comments on prior selection

We attempted to select hyperparameters for the various priors that are suitable for “general application”. Where we use cut-offs, they were chosen generously to allow a wide range of values. For example, the bounds on the migration rate parameters allow them to vary by a factor of 10,000 across the range, and we do not envisage many situations where allowing a wider range would be useful. Our choice for the hyperparameters *c*, *d* on the scale parameters *σ*^2^, *ω*^2^ corresponds to very diffuse prior distributions for both scales, which allow them to take any value dictated by the data. We emphasize that for all examples shown here we used exactly the same parameter settings, and the variety of different EEMS obtained suggests that our priors are sufficiently flexible to be appropriate in a wide range of problems.

## Markov Chain Monte Carlo estimation

We use Markov Chain Monte Carlo (MCMC) to estimate the EEMS parameters by sampling from their posterior distribution given the observed genetic dissimilarities *D*. The two Voronoi tessellations – which describe the spatial organization of the migration rates *m* and the diversity rates *q*, respectively – are independent of each other and are estimated with birth-death MCMC because the number of Voronoi cells is unknown. That is, every move proposes to either add a new cell or remove an existing one (if there are at least two cells). Then the location and rate parameters of a randomly chosen cell are updated with a random-walk Metropolis-Hastings step. (A cell in the migration Voronoi has an effective migration rate, a cell in the diversity Voronoi has an effective diversity rate.) See Supplementary Methods for further details about the computational methods implemented in EEMS.

### Empirical datasets

EEMS is a general method for visualizing spatial population structure and it might be appropriate to apply quality control steps that are customary in population structure analyses, such as pruning SNPs because of long-range LD or high missingness. Measures for SNP and sample quality control have been applied to each of the empirical datasets analyzed here.

#### Elephant data

The African elephant dataset is collected and genotyped as part of a large collaborative study to develop assignment methods for determining the geographic origin of elephant samples from across Sub-Saharan Africa [9]. Samples from both forest and savanna elephants have been collected and genotyped at 16 microsatellite loci. Although the two subspecies can be accurately discriminated using the 16 microsatellites, there is observational and genetic evidence that forest and savanna elephants can hybridize [9]. We analyze the ge-referenced data from [32], which excludes putative hybrids and consists of 211 forest and 913 savanna elephants.

#### Human European and African data

The European dataset was collected and genotyped as part of the POPRES (Population Reference Sample) project [37] and can be accessed at https://www.ebi.ac.uk/ega/studies/phs000145.v2.p2. We used a focal subset of 197,146 autosomal SNPs and 1,379 individuals analyzed in a previous publication [20], with the individual IDs and marker list available from https://github.com/NovembreLab/Novembre_etal_2008_misc. We analyzed a subset of 1,201 individuals from 13 Western European countries: Austria (AT), Belgium (BE), Denmark (DK), France (FR), Germany (DE), Ireland (IE), Italy (IT), Netherlands (NL), Portugal (PT), Scotland (Sct), Spain (ES), Switzerland (CH), United Kingdom (UK). The samples from Switzerland (CH) are split into three subpopulations: French, Italian and German speaking Swiss, coded as CHf, CHi and CHg, respectively. We removed five samples from Italy (7623, 33242, 34049, 38532, 49500) that project outside the main Italian cluster in PC1-PC2 space and therefore are identified as possible outliers in [20]. [For example, these samples might have insular Italian ancestry – Sardinian or Sicilian.] The resulting dataset is described in Supplementary Table 1.

The African dataset was compiled from a subset of two published SNP array datasets: one presented in [41] and available from http://jorde-lab.genetics.utah.edu/?page_id=23 and the other presented in [42] and available from http://www-evo.stanford.edu/repository/. From the Xing *et al.* dataset we extracted the populations: Alur (Al), Bambaran (Ba1), Dogon (Do), Hema (He), Nguni (Ng), Pedi (Pe) and Sotho/Tswana (ST); from the Henn *et al.* dataset we extracted all samples from the populations: Bamoun (Ba2), Brong (Br), Bulala (Bu), Fang (Fa), Hausa (Ha), Igbo (Ig), Kaba (Ka), Kongo (Ko), Mada (Ma2), Mandenka (Ma3) and Xhosa (Xh) and as well as the Yoruba (Yo) samples in the Human Genetic Diversity Project (HGDP) and the Luhya (Lu) and Maasai (Ma1) samples in the HAP1117 subset of HapMap phase 3 [62]. The two subsets were merged at SNPs that have been genotyped in both datasets. From the merged dataset, we then removed SNPs with more than 5% missingness per marker and samples with more than 5% missingness per individual, as well as two Hema individuals that are classified as likely relatives and outliers in most analyses of Sub-Saharan samples in [38]. After these exclusions, we analyzed a dataset composed of 314 samples from 21 Sub-Saharan populations genotyped at 27,825 polymorphic SNPs. The resulting dataset in described in Supplementary Table 2.

#### *Arabidopsis thaliana* data

The *Arabidopsis thaliana* dataset was collected and genotyped as part of the RegMap (Regional Mapping) project [45] and is available at http://bergelson.uchicago.edu/regmap-data/. We downloaded unimputed SNP genotypes for 1,193 samples with high-quality geographic coordinates (latitude and longitude), categorized into twelve geographic regions. From these we analyzed 1,160 accessions from North America and Europe, genotyped at 214,051 SNPs using the Affymetrix Arabidopsis 250K SNP chip [45]. These include 180 accessions from the region *Americas* and 980 accessions from the (European) regions *British-Isles*, *Fennoscandia*, *France*, *Iberia*, *North-West Europe*, *South-Central* and *Austria-Hungary*. We excluded three accessions (Yo-0, Van-0, Buckhorn Pass) from the western coast of North America because the rest are collected from the eastern and central United States, as well as one accession (Can-0) because it is collected from Spain’s Canary Islands and one accession (Da(1)-12) from the Czech Republic because its exact latitude/longitude coordinates are missing.

